# Sparkling Water Limits Cognitive Fatigue with Stable Pupil Diameter during Real-World Esports Training

**DOI:** 10.1101/2025.09.21.677562

**Authors:** Shion Takahashi, Wataru Kosugi, Junichi Kagesawa, Seiichi Mizuno, Takashi Matsui

## Abstract

Prolonged esports play induces cognitive fatigue with reduced executive function and pupil constriction as a neurobiomarker of prefrontal activity across expertise levels. Since this pupil-linked cognitive fatigue is mitigated by sparkling water intake in casual players, we hypothesized that sparkling water prevents cognitive fatigue and sustains esports performance in esports athletes under real training conditions that reflect competitive practice. To test this hypothesis, nineteen hardcore esports players participated in a randomized crossover trial. Each completed two 3-hour sessions of head-to-head virtual football matches while consuming either sparkling water or plain water. Subjective fatigue, enjoyment, and executive function (flanker task) were assessed, and pupil diameter, heart rate, and salivary cortisol were continuously monitored. In-game performance was also recorded. Accuracy declined and pupil constriction progressed under the plain water condition, whereas executive accuracy was preserved and pupil diameter was maintained over the three-hour session in the sparkling water condition. Subjective ratings of fatigue and enjoyment increased similarly in both conditions. Heart rate, including both mean and peak values, and salivary cortisol concentrations showed no significant differences between conditions. In terms of in-game performance, goals, passes, and interceptions did not differ between conditions, whereas the number of fouls was lower and the number of shots was higher in the sparkling water condition. These findings demonstrate that sparkling water mitigates cognitive fatigue and sustains esports performance in esports athletes under real training conditions. As a caffeine- and sugar-free beverage, sparkling water may serve as a practical, low-risk intervention for managing cognitive fatigue and supporting daily practice in modern esports settings.

## 1. Introduction

Prolonged or cognitively demanding activities can lead to a decline in executive function, known as cognitive fatigue, even in individuals with substantial expertise in the task at hand (Sousa et al., 2020; Sun et al., 2022). Unlike traditional physical sports, esports are defined as competitive sports using video games and require high-level cognitive performance rather than physical ability (Bediou et al., 2018). As a result, esports players also experience cognitive fatigue frequently. Previous study showed that both casual and hardcore esports players experience cognitive fatigue after three hours of play (Matsui et al., 2024). Given that cognitive fatigue impairs task efficiency and increases the risk of errors and accidents, it represents a significant issue (Sasahara et al., 2015; Schmidt et al., 2009). Addressing this problem is therefore essential from both health and performance perspectives.

To manage cognitive fatigue, many individuals, including esports players, consume beverages such as energy drinks and sweetened coffee (Alves-Costa et al., 2025; Pang et al., 2025). These drinks contain caffeine and glucose, both of which can enhance executive function after ingestion (Sainz et al., 2020; Wu et al., 2024), and thereby delay the onset of cognitive fatigue (Kennedy & Scholey, 2004). However, excessive or prolonged consumption of caffeine and glucose is associated with a range of adverse health effects, including irregular heart rhythms (Berger & Alford, 2009), sleep disturbances (Machado-Vieira et al., 2001), impaired glucose tolerance (Bedi et al., 2014), and heightened depressive symptoms (Kaur et al., 2020). These potential risks highlight the need for safer and more sustainable beverage alternatives to mitigate cognitive fatigue.

As a potential countermeasure to cognitive fatigue that avoids the health risks associated with caffeine and sugar consumption, sparkling water has recently garnered scientific interest. In addition to enhancing subjective mood, such as increasing feelings of refreshment and reducing subjective fatigue, its consumption helps to maintain fundamental cognitive performance (Fujii et al., 2022; Smit et al., 2004). In our previous study, we found that among casual esports players, consuming sparkling water during a 3-hour gaming session attenuated both cognitive fatigue and the associated reduction in pupil diameter (Takahashi et al., 2025). This effect is thought to be mediated by a neural mechanism in which the carbon dioxide in sparkling water stimulates TRPA1 channels expressed in the trigeminal nerve and TRPV1 channels expressed in the pharyngeal region (Wang et al., 2010; Tsuji et al., 2020). These channels are part of neural pathways that activate the brainstem and its projections to the prefrontal cortex, an area responsible for executive functioning (Tsuji et al., 2020; Zhang et al., 2020). Thus, sparkling water may mitigate cognitive fatigue by indirectly supporting prefrontal cortical activity.

Although the anti-cognitive fatigue effects of sparkling water are demonstrated in casual players, their applicability to esports athletes remains largely untested. To sustain the cognitive performance required during play, hardcore esports players frequently consume large amounts of energy drinks or coffee in their daily routines (Pang et al., 2025). If sparkling water proves effective in this group as well, it could provide a practical beverage strategy independent of caffeine or sugar intake. However, cognitive fatigue manifests differently in casual and esports athletes. In our previous study, extended esports play impaired flanker task performance in divergent ways: casual players displayed prolonged interference times, whereas esports athletes showed reduced accuracy in incongruent trials (Matsui et al., 2024). These differences are likely attributable to task strategies, such as prioritizing speed over accuracy, which may also be linked to distinct patterns of brain activity (Heitz & Schall, 2012; Pastötter et al., 2011). Therefore, whether sparkling water confers comparable anti-cognitive fatigue benefits in esports athletes remains to be determined.

In contrast, both caffeine intake and physical exercise, which are interventions previously demonstrated to enhance cognitive function, activate the prefrontal cortex irrespective of task proficiency (Brunyé et al., 2010; Wu et al., 2024; Yanagisawa et al., 2010; Liu et al., 2024). Similarly, the consumption of sparkling water increases regional cerebral blood flow in the prefrontal cortex, particularly in the orbitofrontal region, and maintains pupil diameter, an indirect marker of activity in the ascending arousal system (Kosugi et al., 2024; Takahashi et al., 2025). These findings suggest that sparkling water may exert ergogenic-like effects on brain function, comparable to caffeine, by facilitating prefrontal activation. Accordingly, sparkling water may help mitigate cognitive fatigue regardless of an individual’s level of expertise.

Therefore, this study aimed to examine whether sparkling water prevents cognitive fatigue and sustains performance in hardcore esports players. In contrast to earlier findings obtained in casual players under standardized laboratory settings, the present study was conducted under real training conditions that closely reflect competitive practice, including head-to-head matches and extended play. This design enhances ecological validity and allows us to test whether sparkling water can be a practical strategy for daily training, where players often rely on caffeinated or sugary beverages despite their health risks. By evaluating its ability to both mitigate cognitive fatigue and maintain performance in authentic training contexts, this study provides new evidence for sparkling water as a sustainable intervention in competitive esports.

## 2. Materials and Methods

### 2.1. Participants

We recruited 20 esports athletes from gaming communities in Akihabara, Japan. They reported regularly practicing virtual football (eFootball, Konami Digital Entertainment Co., Ltd., Japan) for competitive purposes. The definition of a esports athlete was based on the criteria described in Matsui et al. (2024). Of these, 19 participants (23.6 ± 3.6 years old; all male) were included in the final analysis, with one excluded due to data recording errors (Table 1). All participants met the following inclusion criteria: age between 18 and 35 years, ability to understand and provide written informed consent, normal color vision, and availability to attend the experiment in person. Ethical approval for this study was obtained from the Research Ethics Board of the University of Tsukuba (Approval ID: 021-112), and all participants provided written informed consent in accordance with the Declaration of Helsinki.

**Table 1.**
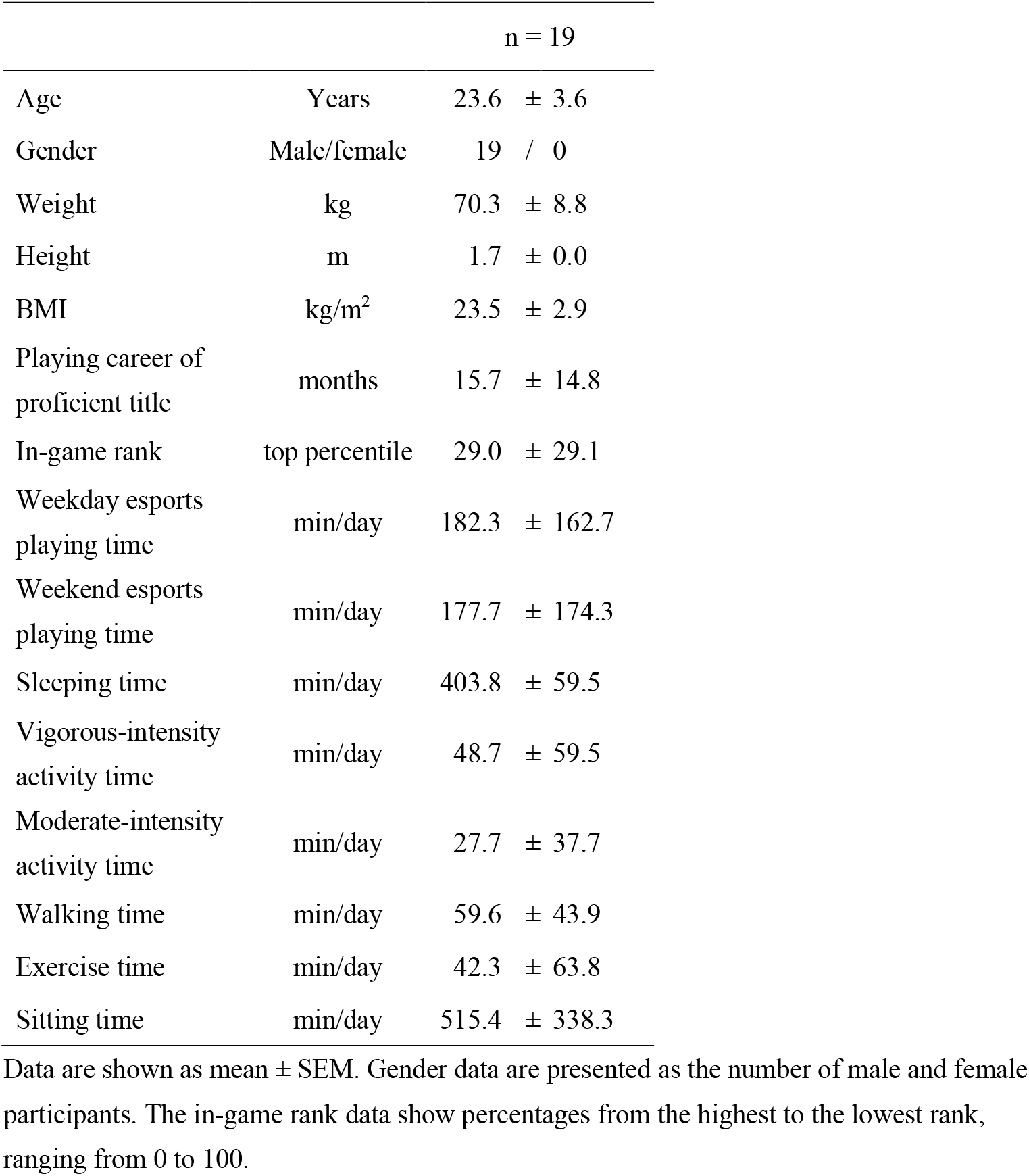
Participant characteristics of playing lifestyles.

### 2.2 Pre-experiment questionnaire

Following the methods outlined by Monma et al., (2024), demographic information, including esports engagement, physical activity, and sleep duration, was collected through a questionnaire administered via Google Forms. Upon recruitment, participants received an email containing a link to the survey and a description of the study, and they were instructed to submit their responses by the morning of the measurement day. Esports involvement was assessed based on participants’ self-reported primary game title, length of experience (in months), competitive rank, and average daily playtime on both weekdays and weekends. Physical activity levels were assessed using the Japanese version of the International Physical Activity Questionnaire (IPAQ), which has been validated by Murase et al. (2002).

### 2.3. Body composition

Body composition was assessed using a bioelectrical impedance analyzer (Seca mBCA 515; Seca, Hamburg, Germany), a device commonly used in esports-related studies (Matsui et al., 2024; Krarup et al., 2024). During the assessment, participants stood barefoot on the device while four electrodes were attached to their limbs. Waist circumference was then measured. Height, weight, body mass index (BMI), fat mass, regional skeletal muscle mass, body water content, and energy expenditure were recorded. All measurements were performed in the morning before the experiment.

### 2.4. Experimental protocol

The experimental procedure followed the method previously validated by Matsui et al. (2024) for assessing cognitive fatigue in hardcore esports players. To examine the effects of beverage intake on cognitive fatigue, a randomized crossover design was employed with two beverage conditions: plain water (PW) and sparkling water (SW), both consumed during play. Participants completed both conditions, with the second session conducted 2 to 30 days after the first, following the same protocol but under the alternate beverage condition. Participants were asked to abstain from alcohol consumption from the day prior to the experiment until its completion, to avoid caffeine intake and strenuous physical activity on the day of the session, and to refrain from eating during the two hours preceding the experiment.

The experimental protocol is illustrated in Figure 1. On the day of data collection, the laboratory environment was controlled at a temperature of 24 ± 2°C and a relative humidity of 50 ± 10%. Upon arrival, participants were verbally informed about the experimental procedures and subsequently provided written informed consent. Prior to the commencement of the experiment, they completed a practice session of the cognitive task under the same environmental conditions as the actual test. A heart rate monitor (Polar H10; Polar, Finland) was then affixed to the chest of each participant, followed by a 10-minute seated rest period.

**Figure 1.**
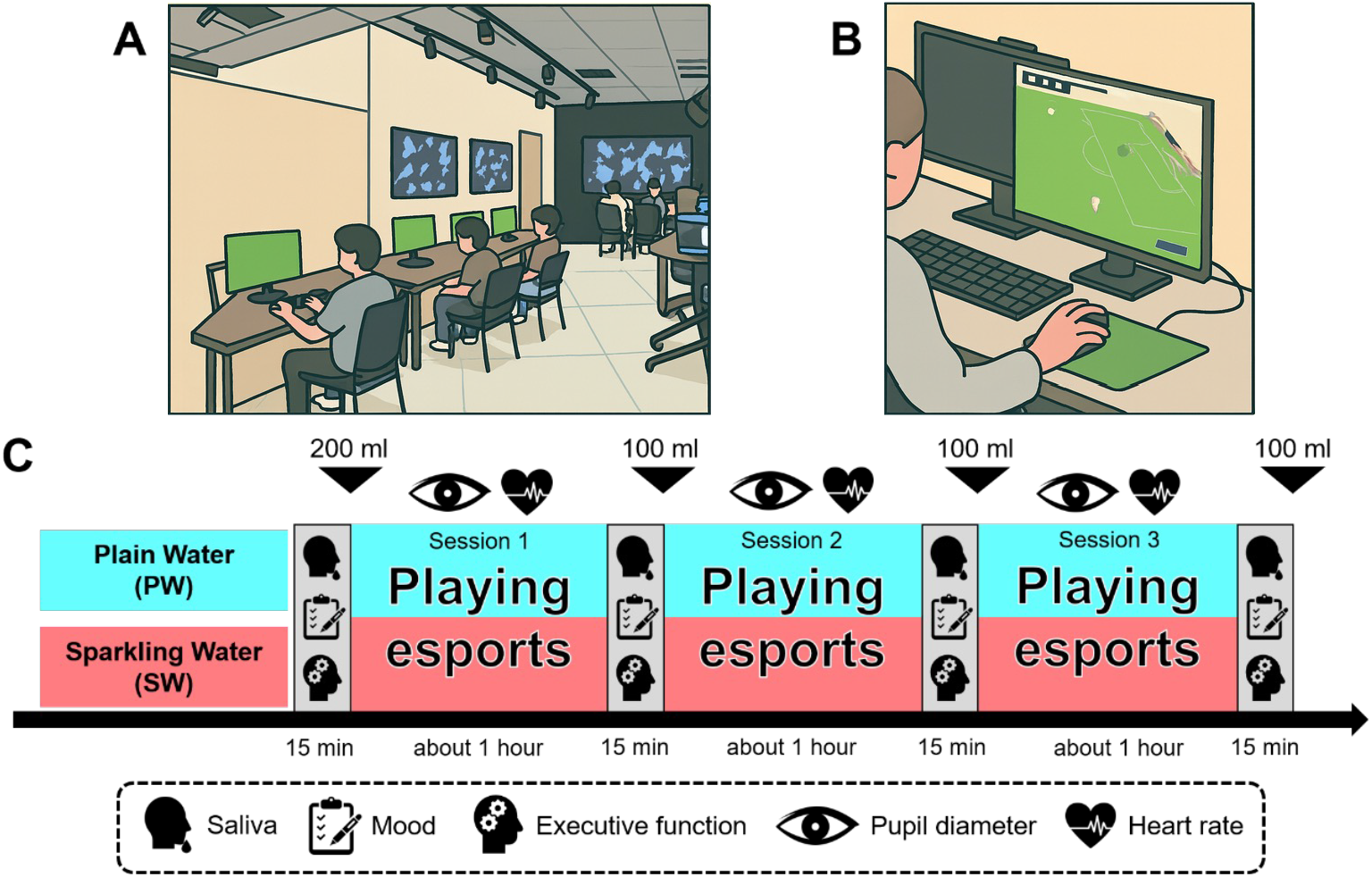
Overview of the experimental protocol under real training conditions. **A**. Esports athletes under real training conditions. **B**. Experimental setup for esports. **C**. Experimental protocol. The same protocol was applied to the plain water (PW) and sparkling water (SW). Participants engaged in eFootball gameplay against other participants for a total duration of three hours. Saliva sample collection, administration of subjective mood questionnaires, and executive function assessments were conducted before the session commenced and at hourly intervals throughout the gameplay. Each assessment session lasted approximately 15 minutes. During gameplay, pupil diameter and heart rate were continuously monitored. At the time points indicated by the downward-facing triangulator in the figure, participants consumed either plain or sparkling water. The beverages were chilled to 4°C, poured into paper cups, and provided to participants immediately before consumption.

To simulate a typical community esports practice session, participants played three consecutive matches against the same opponent. The winner, defined as the player who won the majority of the three games, moved to a different station, while the other remained. However, one participant remained at the same station throughout the experiment in order to allow for continuous measurement of pupil diameter during play.

Before each play session, participants consumed either plain or sparkling water chilled to 4°C, within approximately two minutes. The intake volume was 200 ml prior to the first session and 100 ml before each of the subsequent two sessions, totaling 500 ml. All beverages were refrigerated, opened immediately before consumption, and served in disposable paper cups.

Subjective sensations and executive function were evaluated prior to the first play session (Pre) and at one-hour intervals during the session (1 h, 2 h, and 3 h). Salivary samples were collected at these same time points to assess cortisol levels. Pupil diameter was continuously monitored throughout gameplay using an eye tracker (Tobii Pro Nano; Tobii Technology Co., Ltd., Sweden). Each session lasted approximately 300 minutes, from participant arrival to dismissal. Upon completing the experiment with two conditions, participants received an honorarium of 5,000 yen.

### 2.5. Questionnaires

Participants evaluated two subjective sensations, especially enjoyment and fatigue, using a 100-mm visual analog scale (VAS), following the procedures of previous studies (Tseng et al., 2010; Kotz et al., 2011). The VAS consisted of a horizontal line with the following phrases written at each end: “enjoyed it not at all” to “enjoyed it very much” for enjoyment, and “no fatigue” to “very severe fatigue” for fatigue. Participants indicated their current state by placing a mark along the line. The distance (in millimeters) from the left anchor point was then measured and expressed as a percentage of the total line length.

### 2.6. Executive functions

Executive function was evaluated during and after esports play using a flanker task implemented in PsychoPy3 (v2021.1.2), based on our previous protocol (Matsui et al., 2024). Stimuli were presented in a randomized order, with an equal number of congruent and incongruent trials. Each trial consisted of a fixation cross (250 ms), stimulus presentation (up to 2000 ms), and a blank screen (1000 ms). The stimulus disappeared upon keypress and was followed by a blank screen. Reaction time and response accuracy were recorded. The interference effect was calculated as the difference in reaction time between incongruent and congruent trials.

### 2.7. Pupillometry

Pupil diameter was monitored as an indicator of prefrontal activity during esports play, using an infrared eye tracker (Tobii Pro Nano; Tobii Technology, Sweden), in accordance with the protocol outlined by Matsui et al. (2024). Ambient light levels were controlled and maintained between 250 and 300 lux at the height of the participant’s eyes. The eye tracker was mounted below the display and connected to a dedicated laptop positioned next to the monitor. Participants were seated approximately 60 cm from the display. A four-point calibration was performed before the first session, and pupil data were continuously recorded until the end of the third session. Upon completion of the experiment, only the data corresponding to gameplay periods were extracted for analysis. Pupil diameter was recorded at a sampling rate of 60 Hz, and one-minute average values were computed for further analysis.

### 2.8. Related physiological measurements

Physiological responses to prolonged esports play were assessed using heart rate and salivary cortisol, following previous studies in esports players (Matsui et al., 2024; Cregan et al., 2025). Heart rate was continuously recorded at 1 Hz using a chest-worn monitor (Polar H10; Polar, Finland), which participants wore from the beginning of the experimental session. Salivary cortisol, used as a biomarker of physiological stress, was measured from 2 mL samples collected via straw. Samples were immediately frozen at −80 °C, centrifuged (1,500 × g, 20 min) to remove mucins, and the supernatant was aliquoted and stored at −80 °C. Cortisol concentrations were analyzed in duplicate using a commercial ELISA kit (Salivary Cortisol ELISA Kit; Salimetrics, LLC.) according to the manufacturer’s instructions. Absorbance was measured and concentrations calculated using standard curves on a microplate reader (Varioskan LUX, Thermo Fisher Scientific, MA, USA).

### 2.9. In-game performance

In-game performance was evaluated using statistics displayed on the screen at the end of each match. These statistics were captured using a webcam that directly recorded the monitor on which the participant was playing. For analysis, the average of game statistics across all matches played during each 1-hour session were calculated. Approximately six matches were conducted per session.

### 2.10. Statistical analysis

All statistical analyses were performed using GraphPad Prism version 10 (GraphPad Software, San Diego, CA, USA). Depending on the comparison, data were analyzed using either two-way analysis of variance (ANOVA) or paired t-tests. When significant main effects or interactions were detected, Bonferroni-adjusted post hoc tests were applied. Data are reported as means ± standard error of the mean (SEM), and statistical significance was set at p < 0.05.

## 3. Results

### 3-1. Subjective sensations

Subjective ratings of fatigue and enjoyment, measured using a VAS before and during esports play, showed no significant differences between beverage conditions. Fatigue levels increased over time in both conditions (Figure 2A). Enjoyment ratings were consistently higher during esports play than at baseline, regardless of beverage condition (Figure 2B).

**Figure 2.**
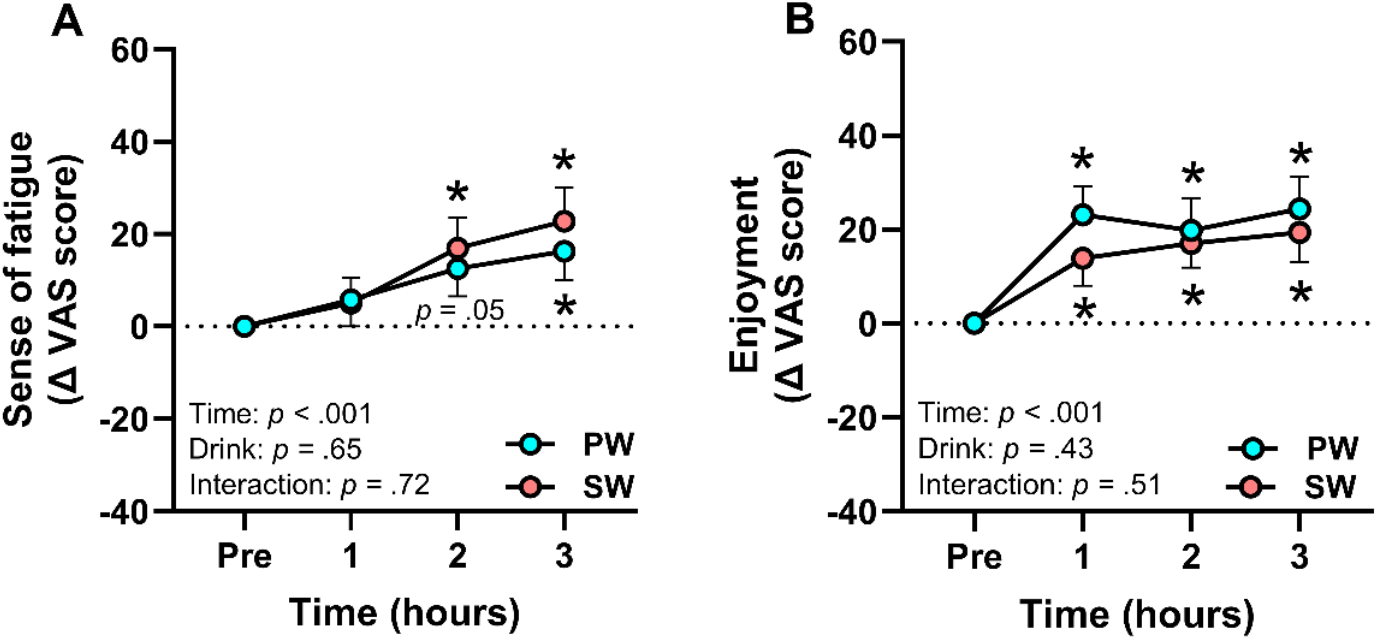
Beverages do not affect subjective sensations during prolonged esports play in esports athletes. **A**. VAS score for the sense of fatigue. **B**. VAS score for the enjoyment. Data are shown as mean ± SEM. Results of two-way ANOVA are shown in the corner of each graph. *Time*: main effect of playing time. *Drink*: main effect of drink condition. *Interaction*: interaction between playing time and drink condition. \**p* <.05 vs. Pre for both groups; *close proximity to a point on the graph. #*p* <.05 vs. FW. The results representing * and # were calculated using Bonferroni’s multiple comparison test.

### 3-2. Executive functions

Flanker interference time remained unchanged throughout prolonged esports gameplay, with no significant effects of beverage condition or time (Figure 3A). In contrast, the correct rate in incongruent trials decreased at the 3-hour time point compared with baseline in the PW condition, whereas accuracy was maintained across the session in the SW condition (Figure 3B).

**Figure 3.**
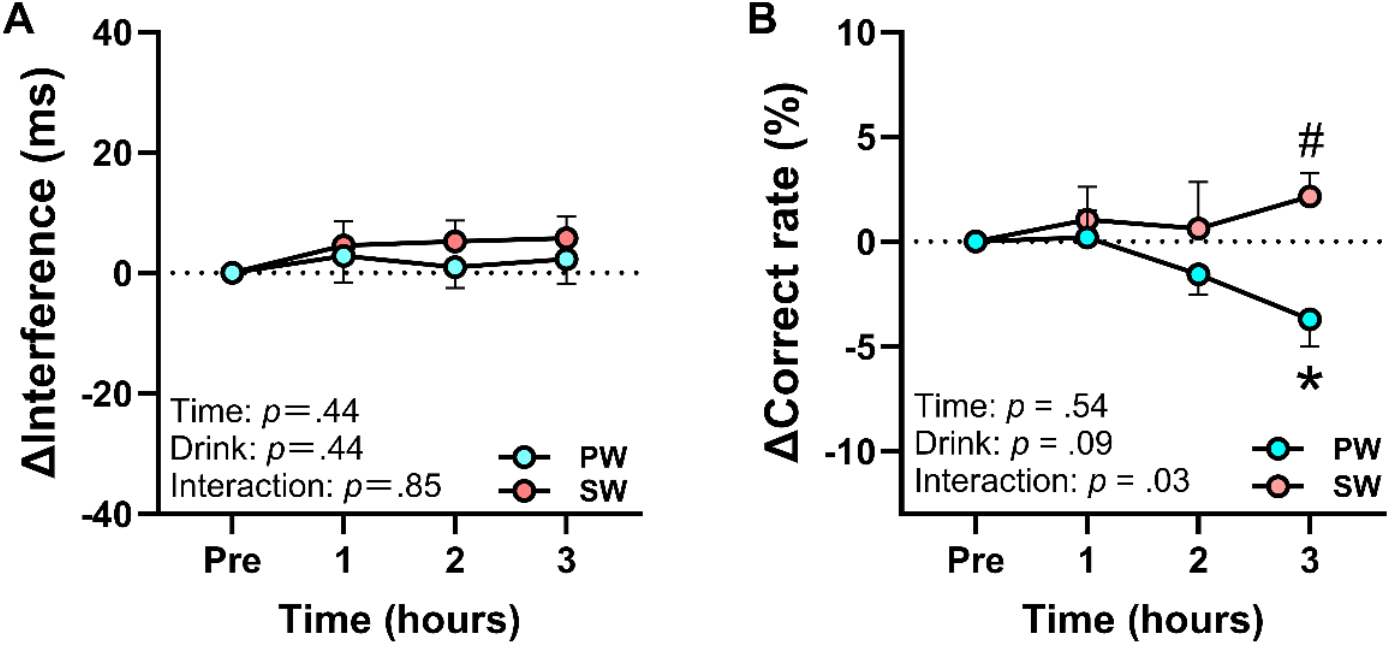
Sparkling water reduces accuracy deterioration associated with prolonged esports play in esports athletes. **A**. Amount of change in the flanker interference time from Pre-point. **B**. Amount of change in the flanker correct rate from Pre-point. Data are shown as mean ± SEM. Results of two-way ANOVA are shown in the corner of each graph. *Time*: main effect of playing time. *Drink*: main effect of drink condition. *Interaction*: interaction between playing time and drink condition. \**p* <.05 vs. Pre for both groups; *close proximity to a point on the graph. #*p* <.05 vs. PW. The results representing * and # were calculated using Bonferroni’s multiple comparison test.

### 3-3. Physiological parameters

In the PW condition, pupil diameter began to decrease at the 2-hour time point and showed a significant reduction at 3 hours. In contrast, no significant change in pupil diameter was observed in the SW condition (Figure 4). No significant effects of time or beverage condition were found for mean heart rate, peak heart rate, or salivary cortisol concentrations across the experimental session (Table 2).

**Table 2.** Physiological parameters of esports athletes during esports play.

|  |  |  | Plain Water |  |  | Sparkling Water |  |  |
| --- | --- | --- | --- | --- | --- | --- | --- | --- |
| Heart rate<br>(bpm) | Resting | - | 81.7 | ± | 3.0 | 81.7 | ± | 2.8 |
|  | Average | 1h | 76.2 | ± | 2.8 | 78.3 | ± | 3.2 |
|  |  | 2h | 73.7 | ± | 2.5 | 74.9 | ± | 2.9 |
|  |  | 3h | 70.9 | ± | 2.0 | 73.2 | ± | 2.5 |
|  | Maximum | 1h | 90.4 | ± | 3.6 | 92.7 | ± | 3.9 |
|  |  | 2h | 87.4 | ± | 3.1 | 88.5 | ± | 3.0 |
|  |  | 3h | 87.4 | ± | 2.7 | 90.0 | ± | 2.7 |
| Salivary cortisol levels<br>(mg/dL) | Resting | - | 0.27 | ± | 0.03 | 0.34 | ± | 0.05 |
|  | Average | 1h | 0.14 | ± | 0.02 | 0.12 | ± | 0.01 |
|  |  | 2h | 0.08 | ± | 0.01 | 0.07 | ± | 0.01 |
|  |  | 3h | 0.07 | ± | 0.01 | 0.09 | ± | 0.01 |

Data are expressed as mean ± SEM. Statistical differences were calculated by Bonferoni's multiple comparisons test. \* $p < .05$ vs. Resting in the group. # $p < .05$ vs. Plain Water group.

**Figure 4.**
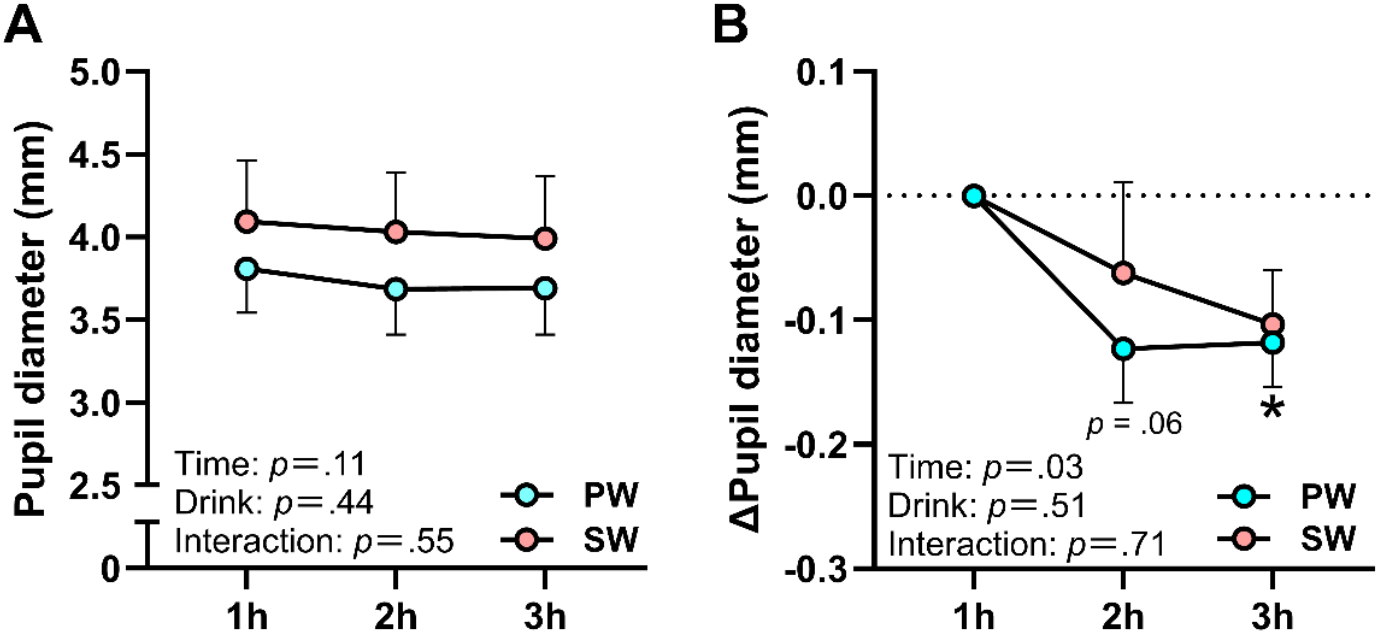
Esports athletes who consumed sparkling water showed no pupil diameter reduction after playing esports for long periods of time. **A**. Pupil diameter during playing esports. **B**. Amount of change in the pupil diameter from 1h-point. Data are shown as mean ± SEM. Results of two-way ANOVA are shown in the corner of each graph. *Time*: main effect of playing time. *Drink*: main effect of drink condition. *Interaction*: interaction between playing time and drink condition. \**p* <.05 vs. Pre for both groups; *close proximity to a point on the graph. #*p* <.05 vs. PW. The results representing * and # were calculated using Bonferroni’s multiple comparison test.

### 3-4. In-game performance

Using the recorded gameplay footage, we computed mean in-game statistics across all matches within each session. To ensure condition-matched comparisons and data quality, analyses focused on participants with complete recordings under both PW and SW (n = 5), as some sessions lacked post-match statistics or contained corrupted files. In this subset, reanalysis of executive function reproduced the overall pattern reported in Section 3.2: flanker task accuracy was higher in the SW condition than in the PW condition (Supplementary Figure 1). Most in-game metrics (points scored, number of passes, and interceptions) were comparable between conditions (Figure 5; Supplementary Figure 2). Notably, Bonferroni-corrected comparisons indicated targeted benefits of SW, with more shots at 3 h and fewer fouls at 2 h relative to PW.

**Figure 5.**
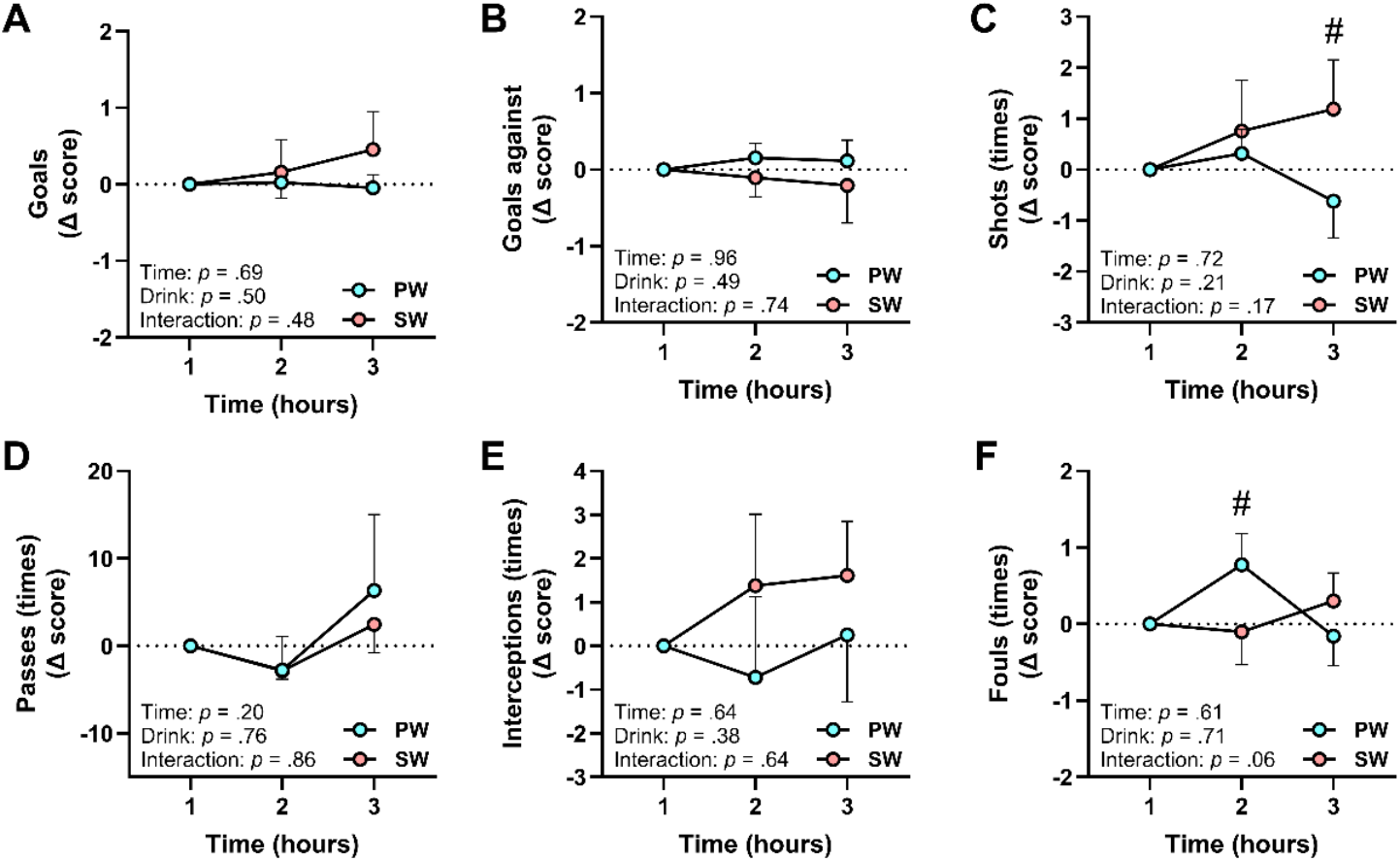
Sparkling water suppresses foul and increases the number of shots in esports athletes. **A**. Goals. **B**. Goals of opponents. **C**. Number of shots. **D**. Number of passes. **E**. Number of interceptions. **F**. Number of fouls. All data are hourly average and the amount of change from 1h. Data are shown as mean ± SEM. Results of two-way ANOVA are shown in the corner of each graph. *Time*: main effect of playing time. *Drink*: main effect of drink condition. *Interaction*: interaction between playing time and drink condition. #*p* <.05 vs. PW. The results representing # was calculated using Bonferroni’s multiple comparison test.

## 4. Discussion

This study demonstrated that sparkling water mitigates cognitive fatigue and sustains cognitive performance in hardcore esports players under real training conditions. In the PW condition, accuracy declined and pupil constriction progressed, whereas in the SW condition, executive accuracy and pupil diameter were preserved. Subjective fatigue, enjoyment, heart rate, and cortisol levels did not differ between conditions. In-game performance analyses showed fewer fouls and more shots in the SW condition. These findings confirm that the beneficial effects of sparkling water are not limited to controlled laboratory settings but are also evident in ecologically valid training environments that reflect the demands of competitive play.

### 4.1. Why sparkling water mitigated cognitive fatigue in esports athletes

In the present experiment, the accuracy of incongruent trials in the flanker task declined at the 3-hour time point under the PW condition (Figure 3B), supporting the findings of our previous study (Matsui et al., 2024). In contrast, sparkling water consumption suppressed both the decline in accuracy and the reduction in pupil diameter observed in the PW condition (Figure 3B, Figure 4). These results are consistent with our prior study using a similar protocol in casual players (Takahashi et al., 2025). Pupil diameter serves as an indirect neurophysiological marker of the ascending arousal system, which is regulated by dopaminergic and noradrenergic activity (Pfeffer et al., 2022). Therefore, the attenuation of cognitive fatigue observed among esports athletes may be attributed to the maintenance of prefrontal cortex activity, facilitated by activation of the ascending arousal system following sparkling water intake.

Unlike the previous study in casual players, where flanker interference time increased with cognitive fatigue (Takahashi et al., 2025), the present study found no significant effects of time or beverage condition on flanker interference time (Figure 3A). This discrepancy suggests that sparkling water does not uniformly enhance executive function but may instead contribute to the maintenance of cognitive endurance over prolonged periods of performance.

Noradrenaline released from the locus coeruleus is known to elevate cortical arousal and strengthen functional connectivity with the salience network, thereby promoting sustained attention (Neal et al., 2023). Furthermore, noradrenaline release from the locus coeruleus is also associated with pupil dilation (Pfeffer et al., 2022). Thus, the suppression of cognitive fatigue and pupil constriction observed in this study may be linked to enhanced cognitive endurance mediated by noradrenaline release, potentially stimulated by sparkling water intake.

### 4.2. Why esports athletes with sparkling water committed fewer fouls and took more shots

In this study, the increase in fouls observed in the PW condition was suppressed when participants consumed sparkling water (Figure 5F). This reduction in aggressive behaviors, such as fouls, is compatible with football analytics showing that disciplinary events accumulate as matches progress in professional leagues, with second-half periods associated with more yellow cards and cautions when demands escalate (R. Sun et al., 2024). Independent evidence indicates that mental fatigue degrades soccer-specific technical and tactical performance, and reviews link mental fatigue to impaired decision quality and execution in match-relevant tasks (Soylu et al., 2022; H. Sun et al., 2022). Against this backdrop, the present pattern is coherent with an interpretation in which sparkling water supports inhibitory control and thereby reduces foul-prone, impulsive actions while promoting fair play. This interpretation is consistent with research connecting executive functions to sport performance and decision-making under pressure, as well as with pupillometry’s role as an index of cognitive control effort related to the locus-coeruleus–norepinephrine system (Vestberg et al., 2012, 2017).

In addition, the number of shots increased in the SW condition (Figure 5C), while shot accuracy (Supplementary Figure 1B–C) remained unchanged. This profile indicates more frequent offensive actions without loss of precision, aligning with evidence that executive functions contribute to efficient action selection and sport-specific decision-making under pressure (Cao et al., 2024). Taken together, these observations raise the possibility that a non-caffeinated, sugar-free beverage strategy could help manage cognitive fatigue and sustain inhibitory control not only in esports training but also in late-match phases of real football, where fatigue-related disciplinary events typically rise (R. Sun et al., 2024).

### 4.3. Why the mood was not changed by sparkling water in esports athletes

In this study, participants’ subjective sensations during prolonged esports play did not differ between beverage conditions (Figure 2). In contrast, our previous study involving casual players found that sparkling water enhanced enjoyment and reduced sense of fatigue (Takahashi et al., 2025). One possible explanation is that the esports athletes in this study regularly engage in extended play sessions of virtual football, the game used in the experiment. As a result, they may not perceive the novelty that casual players experience, and consequently, may not feel any beverage-induced enhancement in enjoyment beyond the baseline of “just playing the game.”

In addition, esports athletes typically train with the goal of winning. In competitive scenarios such as those implemented in the present study, they are likely to mobilize their full cognitive resources, including game-related knowledge and skills. This increased cognitive engagement likely induces a high mental workload (Borghini et al., 2014; Longo et al., 2022). Since mental effort is closely associated with subjective fatigue (Ren et al., 2025), the elevated workload required to perform at their best may have manifested as a stronger sense of fatigue, regardless of beverage condition.

### 4.4 Limitations

This study has several limitations that also point toward directions for future research. First, because carbonation produces distinctive oropharyngeal sensations, full blinding is intrinsically difficult and expectancy effects may influence performance. Future trials should use a taste-, temperature-, and tingling-matched sham beverage under double-blind procedures and pre-specify measurement of expectancy before and after consumption. Placebo and expectancy effects are well documented in exercise and caffeine research and should be explicitly modeled in future analysis plans. Second, we did not directly assay the neural mechanisms through which sparkling water may support executive function. A pragmatic next step is to add lightweight prefrontal fNIRS during training to index cortical activation while maintaining ecological validity, alongside task-embedded pupillometry. fNIRS is feasible in cognitive and sport contexts, and sparkling water has been shown to increase frontal cerebral blood flow relative to still water. In addition, a true dose–response design should vary CO_2_ concentration and bubble characteristics, as bubble size and carbonation parameters modulate oral somatosensory perception and mouthfeel, which likely shape the neural and behavioral responses (Gonzalez Viejo et al., 2019). Third, our sample comprised young male hardcore virtual-football players, which limits generalizability. Cognitive demands differ by esports genre and player profile, with evidence that FPS and MOBA players show distinct attention, inhibition, and decision-making characteristics (Toth et al., 2021; Miao et al., 2024). Multi-genre, multi-cohort replications that include women and broader age ranges are needed to establish external validity.

## 5. Conclusion

Our findings demonstrate that sparkling water sustains cognitive performance in esports athletes under real training conditions by mitigating cognitive fatigue. Executive accuracy and pupil diameter were preserved, and foul counts decreased while shot attempts increased compared with plain water. As a caffeine- and sugar-free beverage, sparkling water can be consumed safely and routinely, making it particularly suitable for daily practice in modern esports settings, where both performance and long-term health are essential. Taken together, these findings highlight sparkling water as a practical, low-risk intervention for managing cognitive fatigue and supporting daily practice in esports and other cognitively demanding activities.

## Declaration of Generative AI in Scientific Writing

During the preparation of this work, generative AI was used for language editing only. After using this tool, the authors reviewed and edited the content as needed and take full responsibility for the content of this publication.

## Data availability

The data supporting the findings of this study are available from the corresponding author upon request.

## Author contribution

W.K., S. M. and T. M. conceived and designed the study. S.T., J.K. and T. M. recruited participants. S.T. and T. M. collected the data, conducted data analysis and interpreted data. S. T. drafted the manuscript. W.K., S. M. and T. M. edited and revised the manuscript. All authors approved the final version.

## Ethics approval and consent to participate

The study protocol was approved by the Research Ethics Board of the University of Tsukuba (Approval ID: 021-112), and all participants provided written informed consent in accordance with the Declaration of Helsinki.

---

**Supplementary figure 1.**
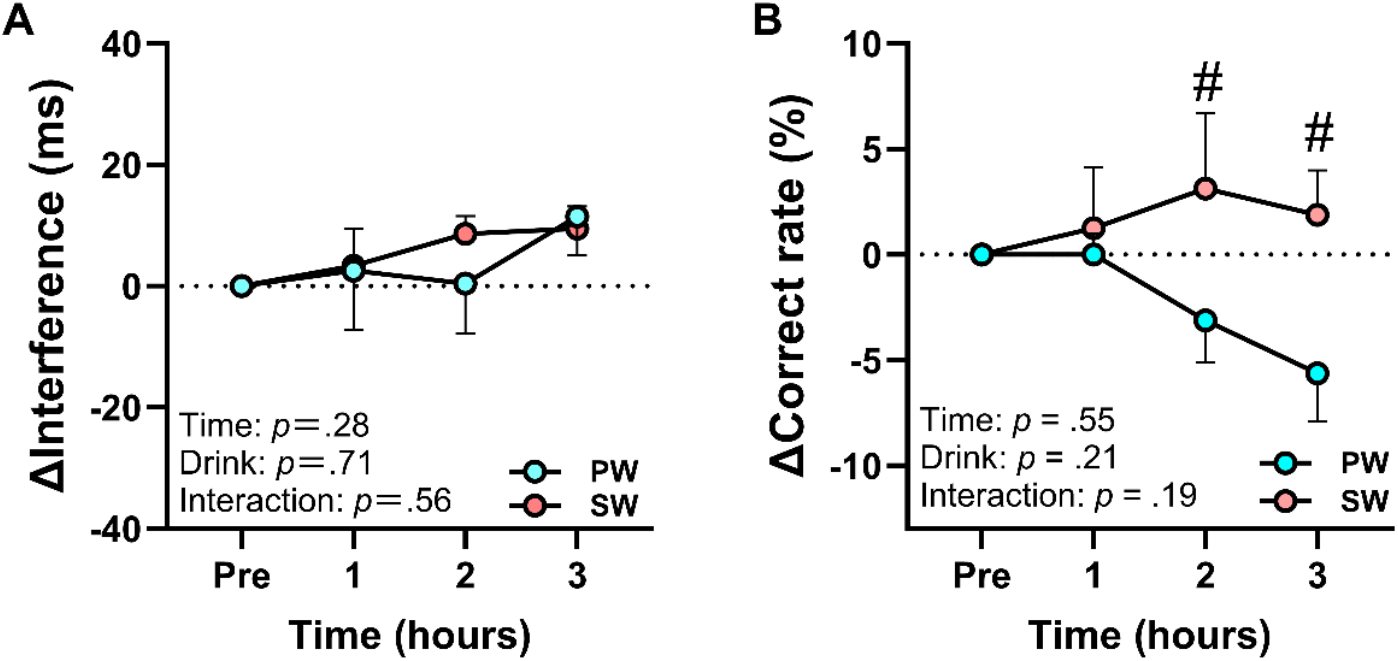
Executive performance of the participants who analyzed game stats. **A**. Amount of change in the flanker interference time from Pre-point. **B**. Amount of change in the flanker correct rate from Pre-point. Data are shown as mean ± SEM. Results of two-way ANOVA are shown in the corner of each graph. *Time*: main effect of playing time. *Drink*: main effect of drink condition. *Interaction*: interaction between playing time and drink condition. #*p* <.05 vs. PW. The results representing # were calculated using Bonferroni’s multiple comparison test.

**Supplementary figure 2.**
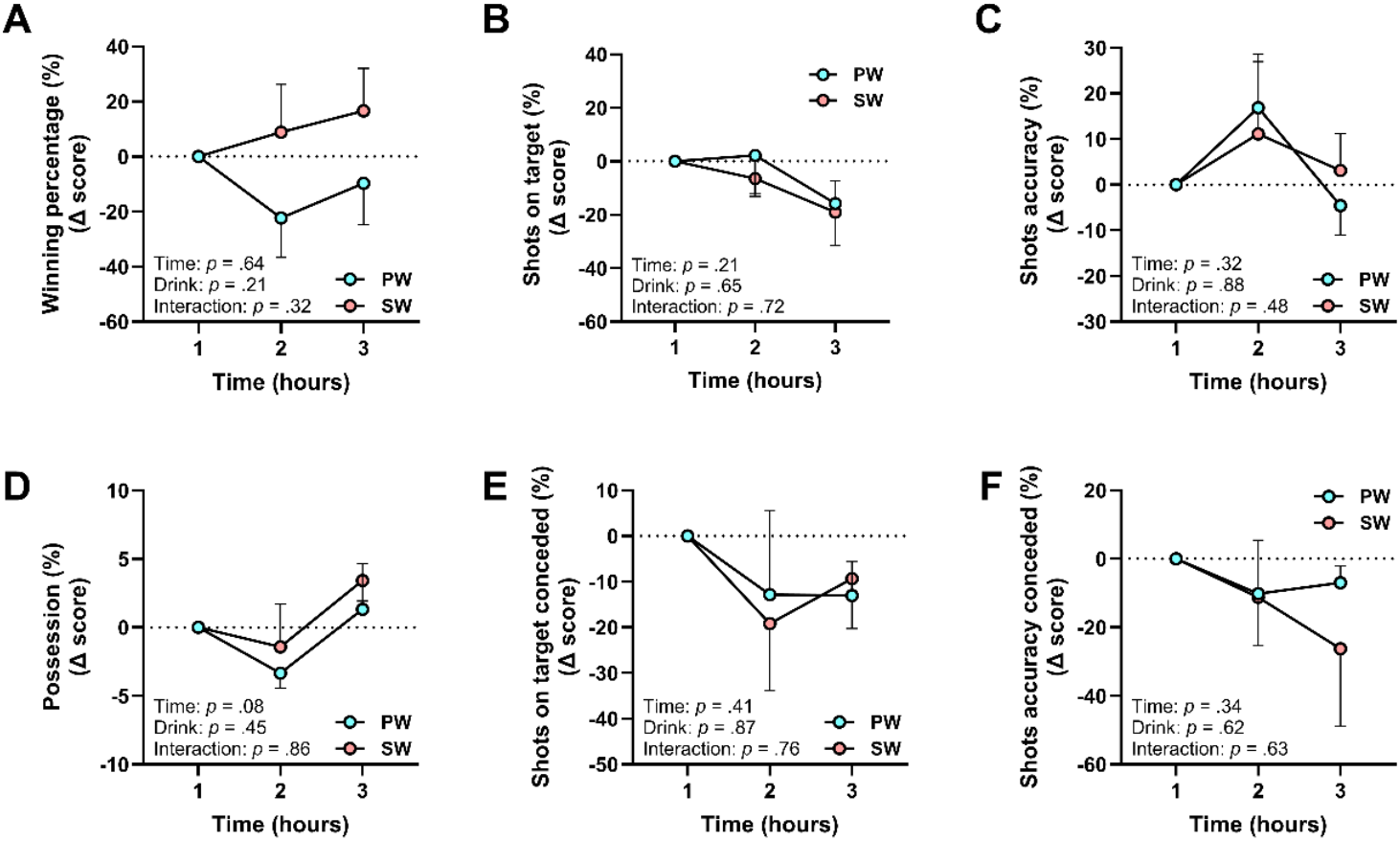
Sparkling water does not adversely affect game performance. **A**. Winning percentage. **B**. Shots on target. **C**. Shots accuracy. **D**. Possession. **E**. Shots on target of opponent. **F**. Shots accuracy of opponent. All data are hourly average and the amount of change from 1h. Data are shown as mean ± SEM. Results of two-way ANOVA are shown in the corner of each graph. *Time*: main effect of playing time. *Drink*: main effect of drink condition. *Interaction*: interaction between playing time and drink condition. \**p* <.05 vs. 1h. The results representing * was calculated using Bonferroni’s multiple comparison test.

